# Fast burst fraction transients convey information independent of the firing rate

**DOI:** 10.1101/2022.10.07.511138

**Authors:** Richard Naud, Xingyun Wang, Zachary Friedenberger, Alexandre Payeur, Jiyun N. Shin, Jean-Claude Béïque, Blake A. Richards, Moritz Drüke, Matthew E. Larkum, Guy Doron

## Abstract

Theories of attention and learning have hypothesized a central role for high-frequency bursting in cognitive functions, but experimental reports of burst-mediated representations *in vivo* have been limited. Here we used a novel demultiplexing approach by considering a conjunctive burst code. We studied this code *in vivo* while animals learned to report direct electrical stimulation of the somatosensory cortex and found two acquired yet independent representations. One code, the event rate, showed a sparse and succint stiumulus representation and a small modulation upon detection errors. The other code, the burst fraction, correlated more globally with stimulation and more promptly responded to detection errors. Bursting modulation was potent and its time course evolved, even in cells that were considered unresponsive based on the firing rate. During the later stages of training, this modulation in bursting happened earlier, gradually aligning temporally with the representation in event rate. The alignment of bursting and event rate modulation sharpened the firing rate response, and was strongly associated behavioral accuracy. Thus a fine-grained separation of spike timing patterns reveals two signals that accompany stimulus representations: an error signal that can be essential to guide learning and a sharpening signal that could implement attention mechanisms.

## Introduction

Optimizing behaviour hinges heavily on adapting internal representations (1, 2). Accordingly, adapting representations is a major focus of machine learning algorithms (3) and neuroscience research (4–7). In the central nervous system, rate codes have been tracked during learning, and two salient attributes have been identified. Firstly, there is *functional reorganization*, whereby training is associated with an increase in the number of task-related neurons (8, 9). Secondly, for neurons already involved in the task, there is a *sharpening* of response selectivity (10). These changes in representations are thought to be mediated by a combination of synaptic plasticity (11), re-structuring local excitation and inhibition (12), and signals feeding back from other brain areas (10, 13).

Theory has hypothesized that two signals accompany these changes in representations. One is a learning signal, a communication that enables efficient coordination of plasticity (14–16). The second is a *sharpening signal*, hypothesized to mediate the sharper responses observed under top-down attention (10, 17). In search for evidence of such signals, we consider that both learning and sharpening have been linked to the phenomenon of bursting (18). For learning, this is because high-frequency firing is a reliable trigger of long-term plasticity in cortex (19–22). For sharpening, this is justified by mechanisms for top-down gain modulation involving burst firing (23–26). Furthermore, bursting is well-situated to communicate such signals as it enables a form of multiplexing (27–30). Previous studies have shown that burst firing conveys information associated with both attention (26, 31, 32) and learning (33, 34), but a reliance on time-averaging prevented investigators to resolve a precise temporal structure. Observing the time course of burst codes is crucial to understanding the interplay between the changes in representations through learning and the occurrence of both learning and sharpening signals.

Here we study the neuronal representations while animals learn to detect an exogenous stimulus: direct electrical micro-ampere stimulation (microstimulation). We have analyzed trial-averaged single-neuron responses to microstimulation so as to obtain a time-resolved picture of bursting modulation (29). Rates of different spike timing patterns were tracked, namely the firing rate, event rate and burst fraction. In general, we found that the timecourse of burst fraction was independent of that of firing rate or event rate. Training had the effect of increasing the number of cells that are responding with an elevated event rate and, in a larger, independent and overlapping group, increased the number of cells responding with increased burst fraction. Concomitantly, we found a delayed error-associated signal that is characterized by an elevated burst fraction. In trained animals, we found that the dynamics of burst fraction responses shifted to earlier times, gradually aligning with the more stable event rate representation. The gradual alignment of burst modulation with the event rate representation sharpened the firing rate representation. Accordingly, the animals’ accuracy was strongly correlated with the early component of the burst fraction. Thus, a burst fraction code resolves error and sharpening signals, highlighting the importance of burst coding for theories of attention and learning.

## Results

We have analyzed juxta-cellular recordings in primary somatosensory cortex (S1) next to a microstimulation (34, 35). In these experiments, animals received microstimulation at random intervals and were rewarded if they licked within 1.2 s of stimulation (Fig. 1A). Any licking after this response window was not rewarded. Juxta-cellular recordings were performed, whereby stereotaxis and spike shape were utilized to help target L5 excitatory cells. We studied the activity of cells in naive animals who have never been exposed to the microstimulation nor its assocation with a reward, and in trained animals that have undergone 1-2 days of training before electrophysiological recording. The same animals returned to the task on multiple subsequent days while their accuracy continued to improve (Fig. 1B).

**Fig. 1.**
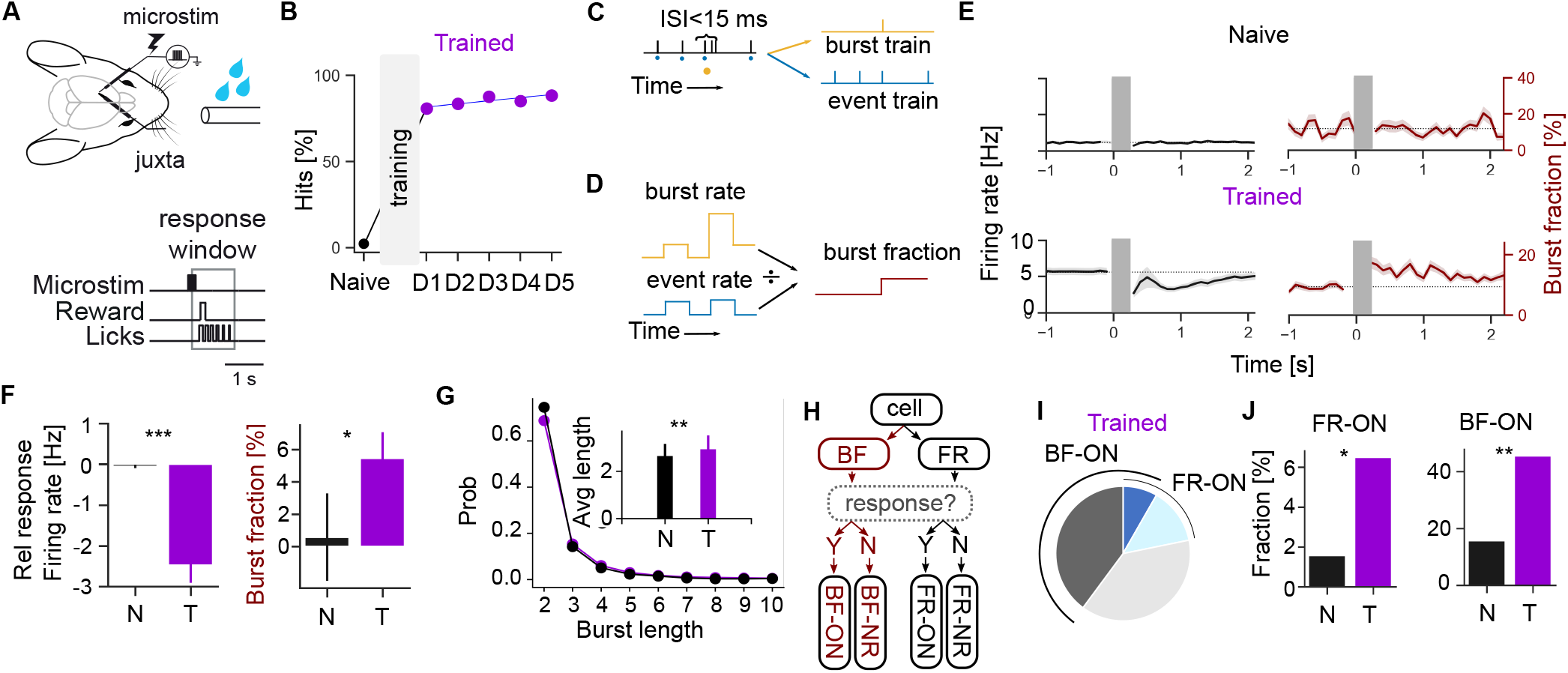
Learning to detect direct electrical microstimulation is associated with the appearance of cells responding with changes with firing rate and cells responding with burst fraction. **A** Schematic of the experimental setup. **B** Progression of average response accuracy through training days, averaged across individual animals. Training lasted 1 to 2 days depending on the animal. After training, Pearson correlation of accuracy vs days after training was 0.86 (*P* =.085, bootstrap test). **C** Schematic of the separation of spike trains into trains of bursts and events based on an interspike interval (ISI) threshold. **D** Schematic of the calculation of burst fraction from the trial averaged burst and event rates. **E** Trial-averaged firing rate (black) and burst fraction (red) averaged across all cells recorded in the naive (top) or trained (bottom) condition. **F** Total response relative to the baseline FR (left) and BF (right) for naive (N) and trained (T) conditions. **G** Histogram of the number of action potentials per bursts in naive (black) and trained (purple) condition. Inset shows the average burst length for naive and trained condition (*P* =.001, rank sum test). **H** Flowchart for the classification of cells according to responses in either FR or BF. Cells showing a significant response in FR are FR-ON cells and cells showing a significant response in BF are BF-ON cells. **I** Pie chart of cell response class in trained animals. **J** The fraction of FR-ON cells (left) and BF-ON cells (right) in naive (N, black bar) and trained (T, purple bar) conditions (FR-ON *P* =.011, BF-ON *P* =.002; proportion z-test). Shaded area along lines in E represents *±* 1 s.e.m. Response to both hits and miss trials are pooled.

These recordings allowed us to unambiguously extract single-neuron responses, except during microstimulation delivery where a stimulus artefact was preponderant. We could unambiguously identify bursts of action potentials (Fig 1C) without confounds induced by spike sorting algorithms (36). Inspired by how synaptic dynamics may read temporal codes (29, 37), we studied burst codes by focusing on trial-averages of different spike timing patterns (Fig 1D): firing rates (FRs) are computed by trial-averages of any spikes, burst rates (BR) are computed by trial averages of bursts, event rates (ERs) are computed by trial-averages of singlets and bursts (i.e. without spikes within a burst). The burst fraction (BF) is computed by dividing the burst rate by the event rate. We focused on ER and BF because they are expected to be modulated independently when bursting arises predominantly from a conjunction of independent signals (5, 29).

To separate the direct effect of stimulation from its learned representation, we first compared responses in naive animals with those in trained animals (Fig. 1). Since electric fields fall as a power law of distance to the stimulating electrode (38), the strength of stimulation is expected to be small at the location of the recording electrode more than 10 *µ*m away. Consistently, neither the average firing rate nor the average burst fraction could reveal a correlate of microstimulation in naive animals (Fig. 1E). In trained animals, the microstimulation caused a decrease of the average firing rate (Fig. 1E-F). This net stimulus-inducced reduction in firing rate does not arise from a global reduction in firing after the microstimulation, as trained animals displayed a class of cells responding with increases in firing rate (FR-ON cells, Fig. 1H). The fraction of FR-ON cells in trained animals, although small, was significantly larger than in naive animals (FR-ON cells, Fig. 1I). These observations recapitulate those of functional reorganization in other animals (8, 9).

Both the burst fraction and the average number of spikes per bursts increased after training (Fig. 1E-G), consistent with previous reports of perceptual learning in auditory cortex (33). Training also increased the fraction of neurons responding with an increase in BF, such that these BF-ON cells (Fig. 1H) were considerably more frequent than FRON cells (*P <*.001, proportion z-test) and more frequent than BF-ON cells in the naive state (*P <*.001, proportion z-test; Fig. 1I). Thus the functional reorganization that is required to bring about the perception of this exogenous stimulus is accompanied with and increase in bursting.

We then asked if BF contained the same information as the FR. The response time-course of single-unit exemplars suggested an independence of FR and BF: while some cells showed a temporally correlated response in FR and BF (Fig. S3), many cells showed non-correlated yet potent responses that were characterized by either a peak time mismatch (Fig. 2), a decreasing FR with increasing BF or a decreasing FR with constant BF (Fig. S3). Ordering cells according to the FR response (Fig. 2B) and showing BF response with the same ordering, we observed that the structure of FR response was not detectable in BF responses (compare Fig. 2B and C). The same holds for comparing ER (Fig. S2) with BF, again suggesting an independence of the representations. Consistently, the time-averaged response magnitude in ER and BF were not correlated across the whole population (Fig. 2D). The firing rate showed a small but significant anti-correlation between FR and BF. These observations are consistent with ER and BF carrying independent signals.

**Fig. 2.**
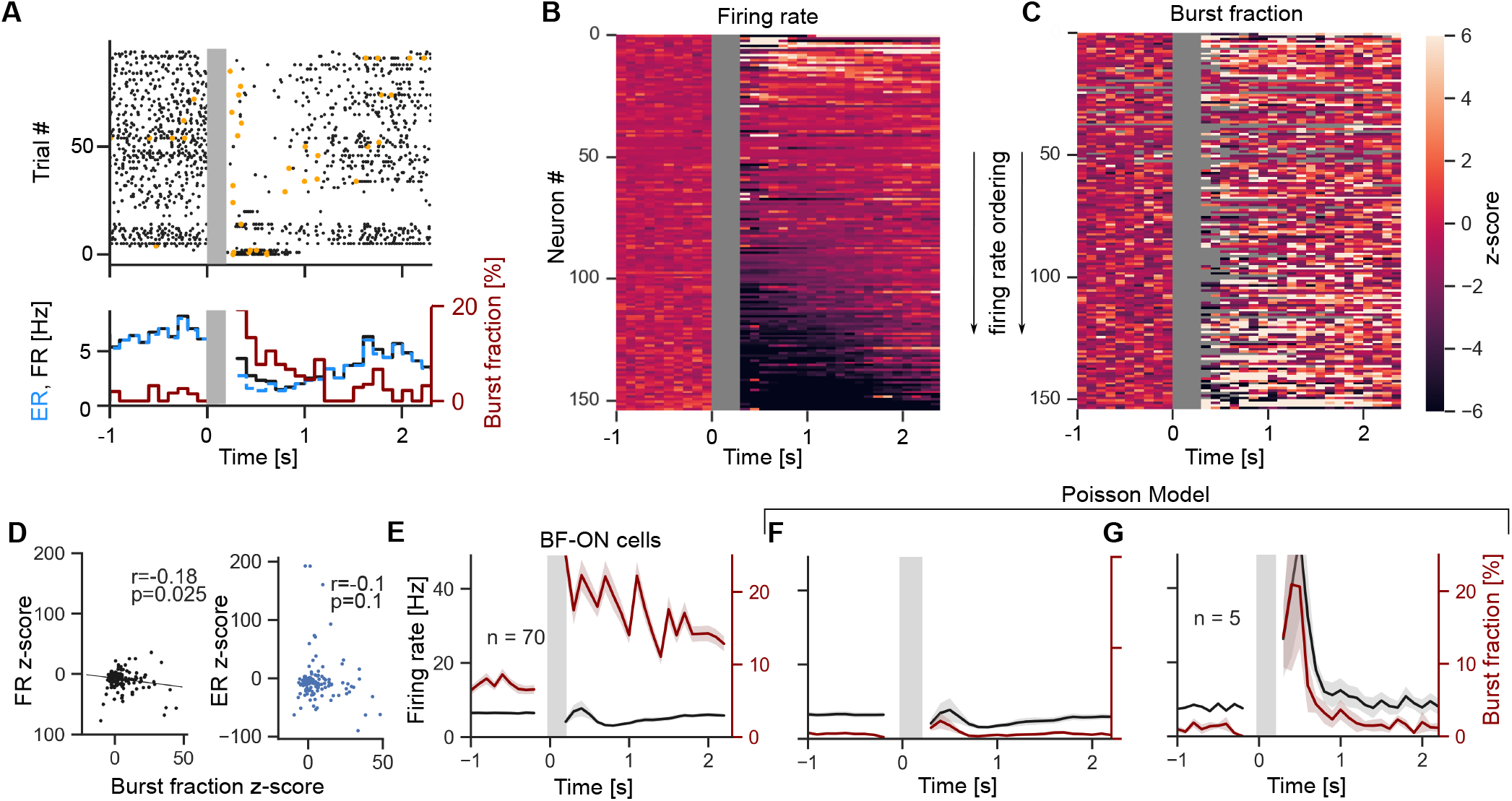
Burst fraction reveals information independent from that in the event rate. **A** For an exemplar cell, the raster of spikes (top, black dots) and first spike in a burst (orange dots) is shown (top). The trial-averaged firing rate (FR, black line), event rate (ER, blue line) and burst fraction (BF, red line, right axis) are shown as a function of time from stimulation (bottom). **B** Heat map of the trial-averaged firing rate response to microstimulation (z-scored) ordered with most firing cells on top. **C** Heat map of the trial-averaged burst fraction (z-scored) keeping the ordering used in B. **D** Scatter plot of post-stimulus z-score of FR against BF and ER against BF. **E** Average FR and BF of all BF-ON cells (*n* =70 out of 153). **F** A Poisson model matching the firing rate of each BF-ON cell recovers the ensemble FR but not the ensemble BF. **G** Average BF-ON cell response from BF-ON cells (*n* =13 out of 279) found by searching through a surrogate Poisson population matching the FR of each cell in the experiment. Shaded area along lines represents *±* 1 s.e.m. Response to both hits and miss trials are pooled.

A widespread abstraction of single-unit response is the inhomogeneous Poisson model, whereby firing is random under specific time-dependent firing rate. This model randomly produces spikes at high frequencies, but the likelihood of these bursts follows the time-dependent fluctuations of the firing rate. To test whether this model can explain the response statistics we observed, we focused on BF-ON cells. Pooling all BF-ON cells revealed a threefold increase of the BF occurring within the 200 ms of stimulation. This fast response relaxed slowly on the time scale of multiple seconds (Fig. 2E). A Poisson model matching the observed FR of each BF-ON cell (Fig. 2F) could not capture these large changes in BF, further indicating that fast burst fraction transients carry information independent of the firing rate. To further test if an inhomogeneous Poisson model combined with possible artefacts arising from performing a selection of responsive cells on rate statistics made of a finite number of trials, we repeated all the data analysis steps on surrogate data generated with a Poisson model that matched the observed firing rates and trial structure (a form of parametric bootstrap, Fig. 2G). In this model and unlike in the real data, only FR-ON cells were selected as BF-ON cells. Accordingly, the model could not account for the long lasting BF increase nor for how widespread BF-ON cells are in the population. Overall, the Poisson model could not account for the independent modulation of BF and FR (black and red curves follow the same time course in Fig. 2F-G). Thus, these observations extend previous reports of FR-independent changes of bursting in monkeys (27, 31, 39) and in rodents (5, 30) and show that the BF responses that arise during training are large and sufficiently fast to show reliable changes within 200 ms.

Theoretical studies have demonstrated that providing neural networks with the ability to independently process sensory information and error signals enables efficient coordination of synaptic plasticity. Specifically, it is the event rate that has been hypothesized to mediate sensory information while the burst fraction was hypothesized to represent error-related signals (15, 40). We thus asked whether and how errors were represented in the population of S1 cells of trained animals. We compared BF, ER and FR in trials where the animal failed to lick in the response window (misses) with trials when the animal was rewarded (hits). While behavioral report of perception is associated with higher levels of activity in higher-order cortex, this is typically not the case for primary cortices (41–43). If a signal is to tell the primary cortex that the current representation gave rise to an erroneous response, it could only come after the time of the perception and the expected time of the response. We thus expected a representation of errors coming after the end of the response window. In some cells (Fig. 3A), we observed increased bursting in misses that occurred late in the trial and was not accompanied with a discernible modulation of the ER. Across all cells, we found that misses were associated with an increase in BF (Fig. 3B) appearing 400 ms after the end of the response window. This error signal was also apparent in the ER and FR, but became significant only later. The timing of these error signals did not match any increase in lick rate (Fig. 3C). Neither licking nor reward were associated with these responses, as seen from catch trials when the stimulation was omitted and when licking was not rewarded (Fig. 3D). Together, we found a delayed representation of error that appeared first in BF but was also detectable in ER and FR.

**Fig. 3.**
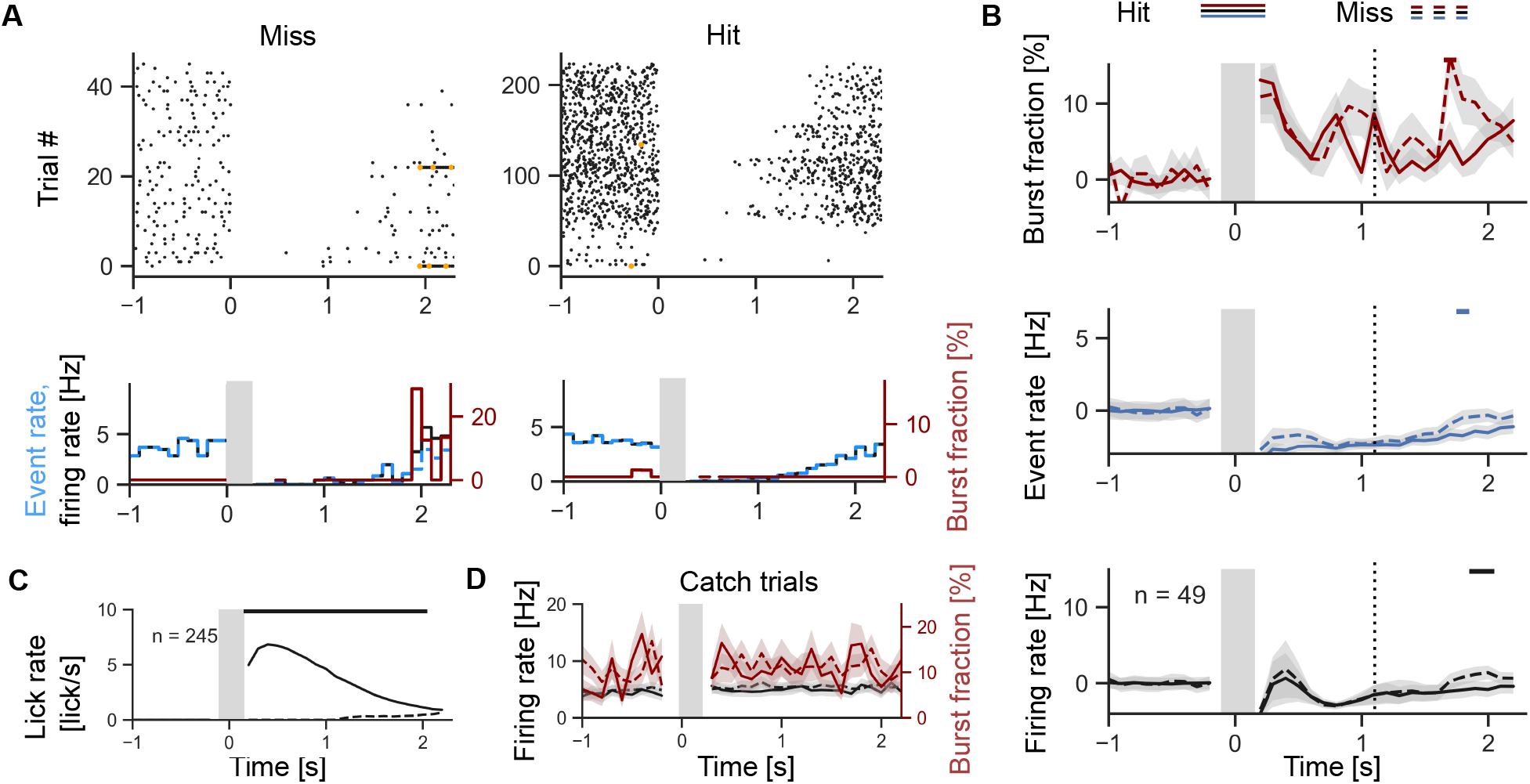
Delayed representation of errors in S1. **A** Raster plot for an exemplar cells (top) separating Hits (right) from misses (left). Bottom panels shows associated trialaveraged firing rate (FR, black lines), event rate (ER, blue lines) and burst fraction (BF, red lines, right axis). **B** Population averaged burst fraction (top), event rate (middle) and firing rate (bottom) for hits (full lines) and misses (dashed lines). The black bar indicates the time bins for which a significant difference was observed (*P <* .05, corrected for multiple comparison at every time bin). The vertical dotted line indicates the end of the response window. **C** Trial and population averaged lick rates for hits (full line) and misses (dashed line). **D** Population averaged firing rate (black) and burst fraction (red, right axis) for catch trials (no stimulation, but rewards are given on licks) separating hits (full lines) from misses (dashed lines). Shaded area along lines represents *±* 1 s.e.m.

In both hits and miss, changes in firing was apparent during the response window. We then focused on hits only, and investigated whether the representations remained stable after the initial 1-2 days of training period. After training, the animals that repeated the task on different days upheld a high accuracy with a non-significant tendency to increase with further days of training (Fig. 2A). Separating the data across later training days, we found that the fraction of FR-ON cells did not show a significant correlation with training days (Spearman correlation *ρ* = 0.2, *P* =.74) nor accuracy (*ρ* = -0.1, *P* =.87), countering the hypothesis that functional reorganization continued at this stage. Turning to the BF representation, we found a similar invariance of the fraction of cells with a significant increase in BF (against days *ρ* = 0.67, *P* =.22, against accuracy *ρ* = 0.81, *P* =.089). Functional reorganization has thus taken place during the first 1-2 days of training and was not noticeable at upon further training.

While the fraction of responsive cells remained constant, it was possible that the amplitude and timing of the response kept evolving. Visual inspection of the responses of all cells suggested a consistent change in the timing of BF response, and a consistent change in the amplitude of the FR responses (Fig. S4). Focusing on cells not showing a firing rate response (FR-NR cells), we found that-even in these cells-the significant BF responses appeared to shift to earlier times as training progressed (Fig. 4A). To quantify the timing changes in either FR, ER or BF, we calculated the responses’ moments (see Methods). We found that the BF moment showed a strong and consistent shift to earlier times on training days when the animals were more accurate (Fig. 4B). Similarly, the FR showed a negative correlation between moment and training days. The same effect, however, was not detectable from the ER. Since the FR (and ER) responses in those cells were always small, it suggests that the timing shift of BF occurs independently of the changes in FR. The shift in BF timing was also significant when considering the whole population (Pearson correlation between BF moment and training days -1.0, *P <* 0.001). During later training, then, it is mainly the timing of bursting modulation which evolved.

**Fig. 4.**
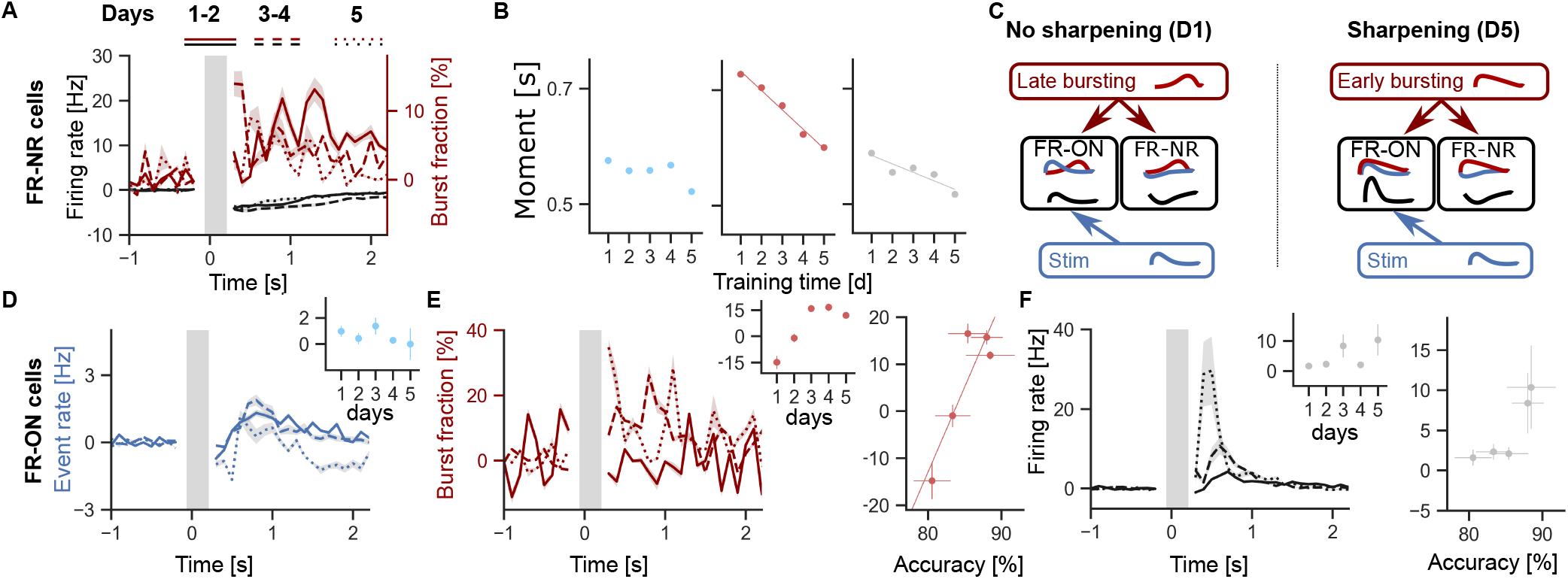
The gradual temporal alignment of a global burst fraction modulation with the focal increase in event rate correlates with behavioral accuracy. **A** Trial- and population-averaged burst fraction and firing rate for FR-NR cells for hits only, separating recordings into subsequent days of recording and training. **B** The firing moment (see Methods) is shown as a function of training days for FR (left), BF (middle) and ER (right), for FR-NR cells. The line indicates a significant correlation (Pearson correlation -0.75 (*P* =.12), -0.99, (*P <*.0001), -0.9 (*P* =.042; bootstrap test), respectively). **C** Schematic: representations progress in later training days mainly by having an earlier modulation of burst fraction. **D-F** Trial- and population-averaged event rate (D), burst fraction (E) and firing rate (F) for FR-ON cells relative to baseline and separated according to training days (legend in panel A). Inset shows relationship between representation in the response period and training days (Pearson correlation -0.59 (*P* = .23), 0.83 (*P* = .84), 0.67 (*P* = .23) for ER, BF and FR, respectively). **E**, right Shows relationship between burst fraction relative to baseline and accuracy. Pearson correlation was significant for the response period (0.89, *P* =.042, bootstrap test). **F**, right Shows relationship between firing rate relative to baseline and accuracy. Pearson correlation was not significant (0.87, *P* =.050, bootstrap test). Responses are for hit trials only.

If the burst modulation of all cells gradually shifts to enter the response window, then this would generate more bursts in cells that were already firing more events while altering minimally the firing rate of the other cells. Such a mechanism can sharpen the selectivity by increasing the response mainly of the cells that were selectively responding to the stimulus and thus made the FR representation more salient (Fig. 4C). To illustrate this, we have separated the responses of FR-ON cells across training days. These cells showed very stable responses in ER (Fig. 4). In addition, these cells had more salient BF responses on days when the animals were more accurate (Fig. 4E). This increased BF multiplies the ER to generate much increased FR responses (Fig. 4F), consistent with previous reports in rodent V1 (12, 44). Thus the sharpening of the selectivity occurred as the timing of burst modulation came for all cells, enhancing the response specifically of responsive cells.

## Discussion

It has long been hypothesized that the distribution of interspike intervals may encode information independent from the FR (45–48), but to read out this information researchers required long time windows and therefore could not reveal time-dependent changes (27, 28, 31, 39) (but see (5)). By exploiting the trial structure, we have shown that the burst fraction reveals a smooth yet at times quickly changing response that is independent of the firing rate. Such reproducible changes in the burstiness have been shown to arise in computational models that provide independent inputs to the apical dendrites (15, 29, 49**?**), a picture that is reinforced by the physiology of hippocampal cells of the CA1 region (5, 50, 51). The burst fraction responses we observed were, however, generally noisier than firing rate responses. Although this may reflect an unavoidable limitation of this neural code, ameliorating the decoding methodologies will certainly improve decoding precision. Some methods have been proposed to address this. Notably, self-tuning algorithms (52) can avoid the reliance on arbitrary parameters (e.g. interval threshold for burst detection). Alternatively, it may be possible to include models of synaptic dynamics in a decoder (29, 37, 53). Furthermore, replacing ensemble averages by weighted readouts that take into account population coding (54) is also likely to improve decoding precision and allow to leverage representation via burst fraction for the decoding of single trials (55) or for establishing communication subspaces (56).

Understanding how the brain learns requires us to establish the precise nature of learning signals and their communication (14). It was hypothesized that changes in BF are well suited for the coordination of learning because they are known to trigger synaptic plasticity (6, 19) and they can be communicated without affecting the processing of sensory information (15). Previous studies have shown that error or reward signals are detectable from field or multi-unit recordings in frontal and sensory cortices (57, 58) with a delay ranging from 100 to 500 ms. We found that the primary sensory cortex represented errors 400 ms after the response window and 1.6 s after the onset of the stimulus. Thus, for these error signals to act on synaptic plasticity, it requires eligibility trace that are of the order of one second (6, 59).

The sharpening of a representation is thought to relate to the difference between responsive and non-responsive cells. During the first stage of learning, a fraction of cells became responsive, but the maximum firing rate of these cells remained small. It was during the second stage of learning that the representation became more salient, increasing the firing rate of responsive cells without changing the firing rate of non-responsive cells. This late stage sharpening of the FR was not due to an increased ER, but rather to an alignment of the burst modulation to the increase in ER. Thus, late-stage learning allowed a temporal alignment of the BF increase with the representation learned in the first stage. Our findings thus implicate BF with both error and sharpening-related information. Correspondingly, top-down modulation is known alter representations with temporal precision (60, 61) and to regulate plasticity (34). Despite the fact that we could not identify the source of the burst modulation, the alignment of BF and ER representation could underlie the construction of ‘vertical assemblies’ (47), that is, burst firing cells arising from the convergence of sensory feedforward and ‘searchlight’ feedback information. Such findings will animate future theoretical work at the intersection between predictive, learning and attention algorithms for adapting representations (15, 16, 62).

## Methods

### Experimental methods

Experimental data has been previously published (34, 35). Briefly, male Wistar rats (Charles River) were housed in reverse 12h light/dark cycle and the experiments were performed during the animal’s night cycle. Experimental procedures were conducted following the guidelines of the Landesamt für Gesundheit und Soziales Berlin.

Before starting head restraint habituation, a metal bolt was implanted on the skull of the animal under ketamine/xylazine anesthesia (100 mg kg ^− 1^ / 5 mg kg ^− 1^). After removing the scalp and periosteum, a thin layer of light-cured adhesive was applied (Optibond, Kerr and Charisma, Kulzer). The head post was glued to the left side of the skull using dental cement (Paladur, Heraeus Kulzer). At least 3 days after the head-post implant, head restraint habituation began. The animals were habituated for 5 min on the first day, and for longer periods the following days until the animal could sit calmly for one hour. Water restriction was established after the second day of habituation (1 mL/day). The animals were then trained to obtain saccharin water (0.1%; Sigma-Aldrich) from the lick port. The weight and general health of the animals was monitored every day. Typical habituation time was 5 days. Craniotomy was performed two days before microstimulation training. A 1.5 mm x 1.5 mm hole was made on the right barrel cortex centered at AP 2.5 mm, ML 5.5 mm from the bregma and a recording chamber was implanted for chronic access. The dura was left intact and the craniotomy was covered with silicon (Kwik-Cast, World Precision Instruments).

Animals were trained to respond to a 200 ms train of microstimulation pulses applied with tungsten microelectrodes (Microprobes) 1500 *µ*m deep in barrel cortex. Stimulation times were random with a refractory period of 1.7 s, corresponding to an exponential inter-trial interval distribution with an average of 17 s. Initial intensity was 160 *µ*A and for the first day, this stimulation was paired with water reward (pairing period). These inter trial statistics were such that only a very small fraction of trials were initiated during the time of the error signal (0.5% of trial had an inter trial interval between 1.7 and 2.1 s). Testing was begun after five pairings. At this stage, animals were only rewarded if they licked within 100 to 1200 ms of the start of the stimulus. Tongue licks were detected using a beam breaker. Licks in the inter-trial intervals were mildly punished by adding 1.5 s to the next stimulus. Once the animals reached 80 % hit rate, the pulse intensity was gradually reduced down to a minimal intensity of 5 *µ*A.

After head-restraint habituation for the naive animals, or after head-restraint habituation and training for the trained animals, juxta-cellular recordings were performed from deep layer neurons in S1. Animals were awake and behaving during the recordings. Glass pipettes (4-8 MO) were filled with extracellular solution containing 135 mM NaCl, 5.4 mM KCl, 1.0 mM MgCl_2_,1.8 mM CaCl_2_, and 5 mM HEPES (7.2 pH). Juxta-cellular signals were amplified and low-pass filtered at 3 kHz with a patch-clamp amplifier (NPI) and sampled at 25 kHz by a Power 1401 data acquisition interface which was itself controlled by Spike2 software (CED).

### Data analysis methods

Recorded neurons were separated into putative fast-spiking (FS) interneurons and regular spiking (RS) pyramidal neurons based on the spike halfwidth and firing rate. Cells with spike half-width of less than 0.5 ms and firing rate higher than 8 Hz were classified as FS. Only RS were used in this study. Also, only cells responding that could be tracked for at least 40 repetitions were included.

Spike trains were parsed in two types of events: isolated spikes preceded and followed by interspike interval larger than 15 ms, and bursts preceded with an interval larger than but followed by one or multiple intervals smaller than 15 ms. Trial-averaging of all spikes was perform to yield FRs. Trial averaging of singlets gave singlet rate (SR) and trial averaging of burst gave the burst rate (BR). We defined the event rate as ER = SR+BR and the burst fraction as BR/ER ((29)). All trial averaging were calculated based on bins of 100 ms centred on the time of the stimulation. Bins were non overlapping and no smoothing was performed in order to preserve the independence of each trial averaged measure. For bins with an event rate of zero, the burst fraction is undefined and this bin is excluded from the following population- or time-averages. It is possible to replace bins were ER is zero by smoothed version of the ER, but we did not do so here in order to ensure statistical independence of each time bin. The 1 s period before stimulation was used to calculate the stationary BF, ER and FR (BF_0_, ER_0_ and FR_0_, respectively) as well as their standard deviation (*σ* _BF_, *σ*_*ER*_ and *σ*_FR_)which were used to calculate the z-scores (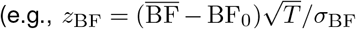, where *T* is the number of time bins in the temporal average 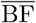 taken after stimulation). Classification of FR-ON and BF-ON cells was based on a z-score threshold of 3 (corresponding to *P* =0.001 for a two-tailed test z-test) for response to hit trials during the period 0.2 to 2.2 s poststimulus.

For Fig. 2F, an inhomogeneous Poisson model was simulated for each BF-ON cell with intensity matching the trial-averaged FR in bins of 100 ms. A FR of 0 was artificially imposed during the microstimulation. The average trial-average FR and BF of the Poisson model was then contrasted with observed quantities. For Fig. 2G, an inhomogeneous Poisson model was simulated for *each* cell in the experiment, with intensity matching the trial-averaged FR and number of trials matching those of experiments. We then searched for BF-ON cells in the simulated response and averaged the FR and BF of these cells.

Moment was calculated according to the formula 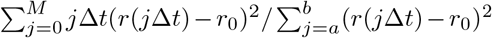 where *r* is the BF for the BF moment, and *r* is the FR for the FR moment. The integration limit *M* was fixed to 2.0s post stimulus ((*a, b*)= (2, 20), Δ.*t* = 100 ms).

Bootstrapped estimates of the p-value of correlations was performed by first computing the correlation of the data. We then shuffled the allocation between recorded cell and recorded day and repeated the calculation of the correlation. We repeated the shuffling 50000 times in order to build the null probability distribution. The p-value was calculated by taking the sum of the probability distribution associated with a correlation whose absolute value is greater than the observed correlation (two-sided test).

