## Supplementary Figures for "Fast burst fraction transients convey information independent of the firing rate"

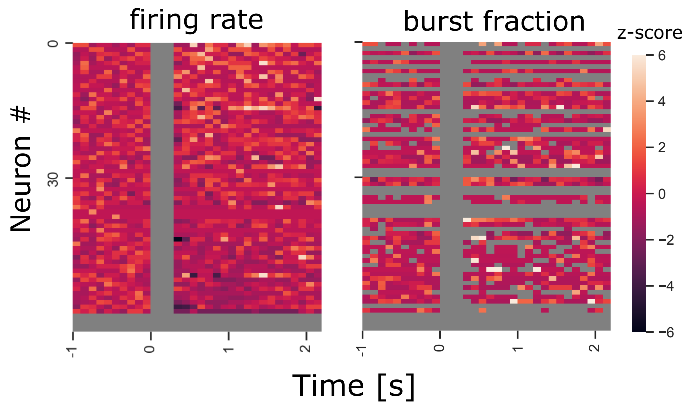


**Fig. S1.** Heat map of event rate across all recorded cells, ordered by average post-stimulus ER response.

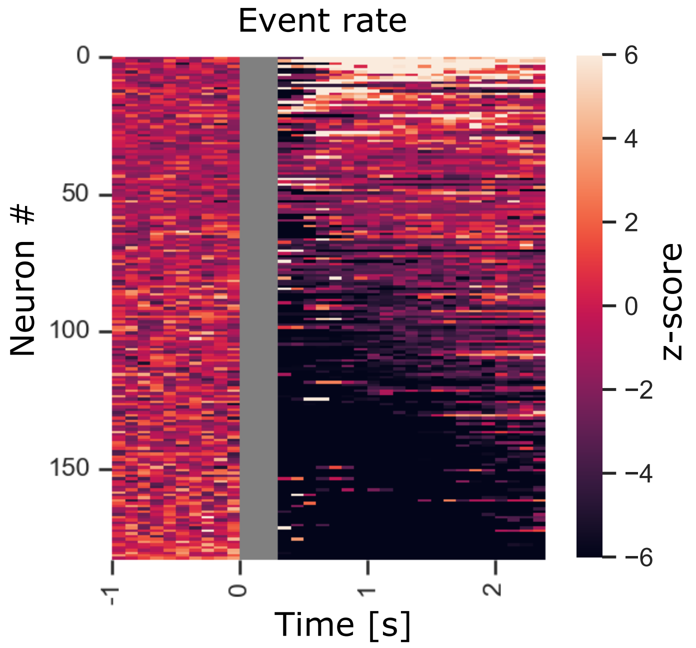


**Fig. S2.** Heat map of event rate across all recorded cells, ordered by average post-stimulus ER response.

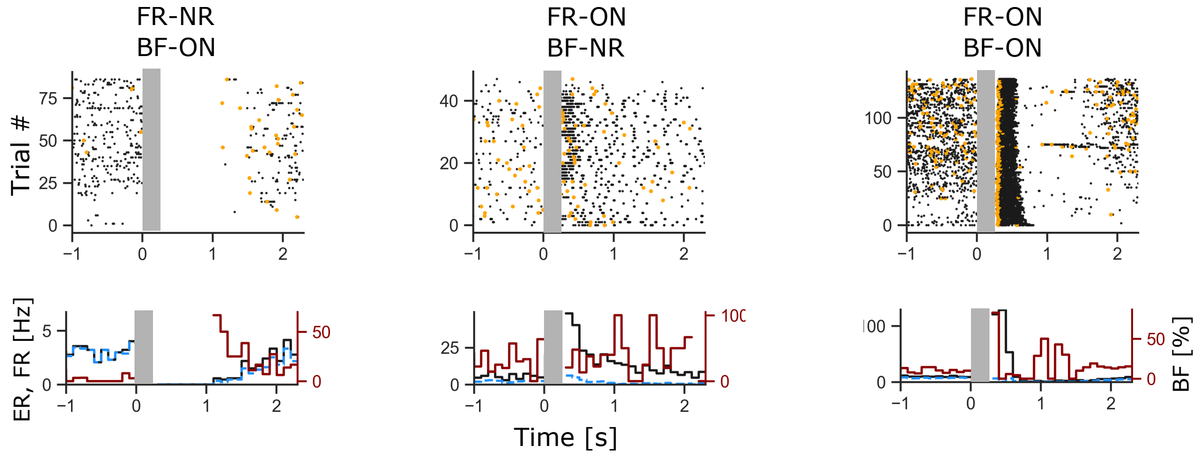


**Fig. S3.** Three exemplars of single cell response to microstimulation (left).

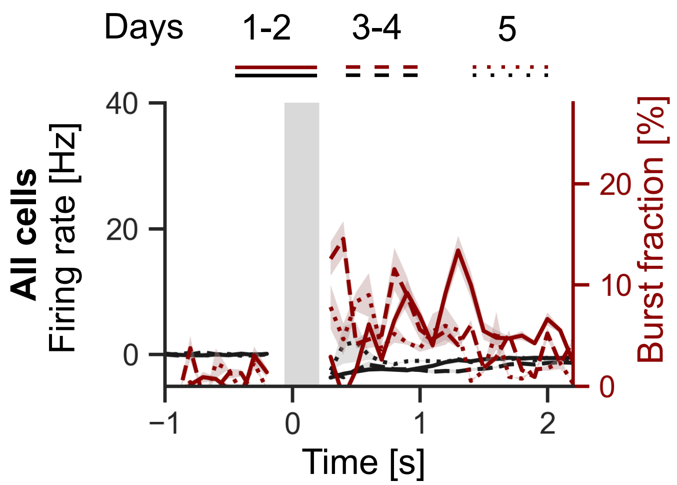


**Fig. S4.** Response in firing rate (black lines, left axis) and burst fraction (red lines, right axis) separated across days for all cells.
